# Using recurrent neural networks to detect supernumerary chromosomes in fungal strains causing blast diseases

**DOI:** 10.1101/2023.09.17.558148

**Authors:** Nikesh Gyawali, Yangfan Hao, Guifang Lin, Jun Huang, Ravi Bika, Lidia Calderon Daza, Hunkun Zheng, Giovana Cruppe, Doina Caragea, David Cook, Barbara Valent, Sanzhen liu

## Abstract

The genomes of the fungus *Magnaporthe oryzae* that causes blast diseases on diverse grass species, including major crop plants, have indispensable core-chromosomes and may contain one or more additional supernumerary chromosomes, also known as mini-chromosomes. The mini-chromosome is speculated to play a role in fungal biology, provide effector gene mobility, and may transfer between strains. To understand and study the biological function of mini-chromosomes, it is crucial to be able to identify whether a given strain of *M. oryzae* possesses a mini-chromosome. In this study, we applied recurrent neural network models, more specifically, Bidirectional Long Short-Term Models (Bi-LSTM), for classifying DNA sequences as core-or mini-chromosomes. The models were trained with sequences from multiple available core- and mini-chromosome assemblies. The trained model was then used to predict the presence of the mini-chromosome in a global collection of *M. oryzae* isolates using short-read DNA sequences. The model predicted that the mini-chromosome was prevalent in *M. oryzae* isolates, including those isolated from rice, wheat, Lolium and many other grass species. Interestingly, 23 recent wheat strains collected since 2005 all carried the mini-chromosome, but none of nine early strains collected before 1991 had the mini-chromosome, indicating the preferential selection for strains carrying the mini-chromosome in recent years. Based on the limited sample size, we found the presence of the mini-chromosome in isolates of pathotype *Eleusine* was not as high as isolates of other pathotypes. The deep learning model was also used to identify assembled sequence contigs that were derived from the mini-chromosome and partial regions on core-chromosomes potentially translocated from a mini-chromosome. In summary, our study has developed a reliable method for categorizing DNA sequences and showcases an application of recurrent neural networks in the field of predictive genomics.

## INTRODUCTION

The fungus *Magnaporthe oryzae*, also known as *Pyricularia oryzae*, is responsible for causing blast diseases on diverse grass species, including major crops (Gladieux *et al*., 2018; Valent *et al*., 2020; Ristaino *et al*., 2021). Rice blast disease, caused by *M. oryzae Oryza* (MoO) pathotype, poses a significant threat to rice production (Fernandez & Orth, 2018). Wheat blast disease, caused by a distinct pathotype, *M. oryzae Triticum* (MoT), emerged in Brazil in 1985 and spread within South America and recently to South Asia and Africa (Malaker *et al*., 2016; Islam *et al*., 2016; Tembo *et al*., 2020). Additionally, blast diseases caused by the *Setaria* pathotype (*M. oryzae Setaria*, MoS) on foxtail millet and by the *Eleusine* pathotype (*M. oryzae Eleusine*, MoE) on finger millet are significant diseases of these ancient subsistence crops (Lenne *et al*., 2007; Sharma *et al*., 2014). Since the early 1990’s, the *Lolium* pathotype (*M. oryzae Lolium*, MoL) has emerged in the US to cause serious blast diseases on popular turf grass or forage crops, including perennial ryegrass, annual ryegrass, and tall fescue. There are other *M. oryzae* strains from other hosts, such as oats (*Avena*), buffelgrass (*Cenchrus*), crabgrass (*Digitaria*), and signalgrass (*Urochloa*), respectively (Gladieux *et al*., 2018; Rahnama *et al*., 2021a).

The genome of *M. oryzae* contains seven essential core-chromosomes and many genomes possess one or a few extra, non-essential supernumerary chromosomes (Dean *et al*., 2005; Peng *et al*., 2019). Besides *M. oryzae*, many plants, animals, and other fungi carry supernumerary chromosomes, which are also known as extra chromosomes, dispensable chromosomes, accessory chromosomes, or B-chromosomes. Supernumerary chromosomes are hypothesized to be an accelerator for fungal adaptive evolution (Croll *et al*., 2013). Supernumerary chromosomes in *M. oryzae* are referred to as mini-chromosomes because their sizes are typically smaller than core-chromosomes (Orbach *et al*., 1996; Peng *et al*., 2019; Langner *et al*., 2021). As compared to core-chromosomes, mini-chromosomes in *M. oryzae* are more repetitive, containing more transposable elements and fewer genes. The repeat-rich characteristic provides ample intrachromosomal homology for DNA duplication, loss, and rearrangements, creating conducive environments to accelerate genome evolution (Peng *et al*., 2019; Huang *et al*., 2023). Indeed, mini-chromosomes are highly variable among *M. oryzae* strains (Peng *et al*., 2019; Langner *et al*., 2021; Liu *et al*., 2022). Mini-chromosomes carry effector genes that can be found in core-chromosomes in different strains, suggesting crosstalk between mini- and core-chromosomes (Chuma *et al*., 2011; Peng *et al*., 2019; Langner *et al*., 2021). Therefore, the mini-chromosome is thought to be capable of mediating the mobility of effector genes, facilitating fungal adaptation.

To confirm and further understand the evolutionary role of the mini-chromosome, it is critical to be able to determine if a *M. oryzae* strain carries the mini-chromosome. Contour-clamped homogeneous electric field (CHEF) electrophoresis of intact chromosomes is the means to provide conclusive evidence for the presence or absence of chromosomes with sizes smaller than core-chromosomes (Orbach *et al*., 1996). However, the technique is laborious and requires specific equipment. This is a significant hurdle as research labs may not have access to all strains published in the literature, especially given that wheat blast causing strains are quarantined in the United States. A reliable method to determine the presence of a mini-chromosome from publicly available or newly generated sequencing data would be fast, cost-effective, and decentralized. A simple strategy is to align sequencing reads to known mini-chromosome genomes for the determination of the proportion of mini-chromosome genomes supported by reads. The lack of knowledge about critical elements required for mini-chromosomes, the high-level of variability among mini-chromosomes, and the potential exchanges between core- and mini-chromosomes complicate the analysis. Fortunately, multiple complete core-chromosome and mini-chromosome genomes are currently available, providing the opportunity to deploy deep learning algorithms to learn features of core- and mini-chromosome sequences for prediction.

Use of neural network based deep learning techniques has rapidly increased due to availability of large data, and their ability to find complex patterns. Recurrent Neural Network (RNN) and, specifically, Long Short-Term Memory (LSTM) networks have been used in genomics to utilize the sequential property of DNA sequences for making various predictions (Shen *et al*., 2018; Liu *et al*., 2019; Zhang *et al*., 2020). An RNN represents a group of artificial neural networks that incorporate feedback connections to retain and utilize information from prior input events as activation. These networks leverage their internal state to process input sequences of varying lengths. However, training RNNs to effectively capture long-term dependencies poses challenges as the error signals flowing backward often suffer from issues of either explosive amplification or rapid attenuation, a.k.a. exploding or vanishing gradients (Bengio *et al*., 1994; Hochreiter & Schmidhuber, 1997). To address this problem, the LSTM architecture was introduced as an extension to the vanilla RNN, a simple form of RNN (Hochreiter & Schmidhuber, 1997). Another enhancement is the Bidirectional LSTM (Bi-LSTM), which considers sequential context from both directions and can improve performance (Schuster & Paliwal, 1997; Baldi *et al*., 1999). In our study, we apply Bi-LSTM deep learning to predict the presence of mini-chromosome sequences based on the genomic sequence data. Experimental results show that a Bi-LSTM neural network model can accurately infer the presence of the mini-chromosome in strains of *M. oryzae*.

## RESULTS

### Training data to assign DNA sequences to core-or mini-chromosomes

The goal of the study was to predict the presence of mini-chromosomes using its genomic sequencing data. Although genomic data of hundreds of *M. oryzae* strains are publicly available, there is very little data regarding if an individual strain possesses mini-chromosome(s). We addressed this by building a Bi-LSTM model to classify DNA sequences as originating from core-or mini-chromosomes (**Figure 1**). The output of the model is used to infer the presence of mini-chromosomes in a strain based on the proportion of mini-chromosome-derived sequences among the total short DNA sequences examined. We collected finished genome assemblies of *M. oryzae* strains with or without the mini-chromosome for model training, from which short sequences were extracted. The strains harboring at least one mini-chromosome include B71 (MoT) (Peng *et al*., 2019), LpKY97 (MoL) (Rahnama *et al*., 2020), TF05 (MoL), and O135 (MoO) (Orbach *et al*., 1996), while the strains containing no mini-chromosomes include the MoO reference strain 70-15 (Dean *et al*., 2005) and MZ5-1-6 (MoE) (Gómez Luciano *et al*., 2019). Approximately 11.2 Mb and 252.2 Mb from mini- and core-chromosomes were collected for model training (**Table 1**). Note that the presence of at least a mini-chromosome in B71 and O135 was verified by CHEF (Orbach *et al*., 1996; Peng *et al*., 2019; Rowe *et al*., 2023). The collected six mini-chromosomes and 42 core-chromosomes were chopped into short DNA sequences and labeled with either mini or core as the sequence source for model training.

**Table 1.** Summary of genome assembly data for model training.

| strain | pathotype | mini (bp) | # mini | core (bp) | # core |
| --- | --- | --- | --- | --- | --- |
| 70-15 | MoO | 0 | 0 | 41,027,733 | 7 |
| MZ5-1-6 | MoE | 0 | 0 | 42,703,282 | 7 |
| O135 | MoO | 1,747,687 | 1 | 41,933,874 | 7 |
| B71 | MoT | 1,903,245 | 1 | 42,908,164 | 7 |
| LpKY97 | MoL | 3,910,017 | 2 | 41,702,954 | 7 |
| TF05 | MoL | 3,598,139 | 2 | 41,915,541 | 7 |
| **Total** | - | **11,159,088** | **6** | **252,191,548** | **42** |

**Figure 1.**
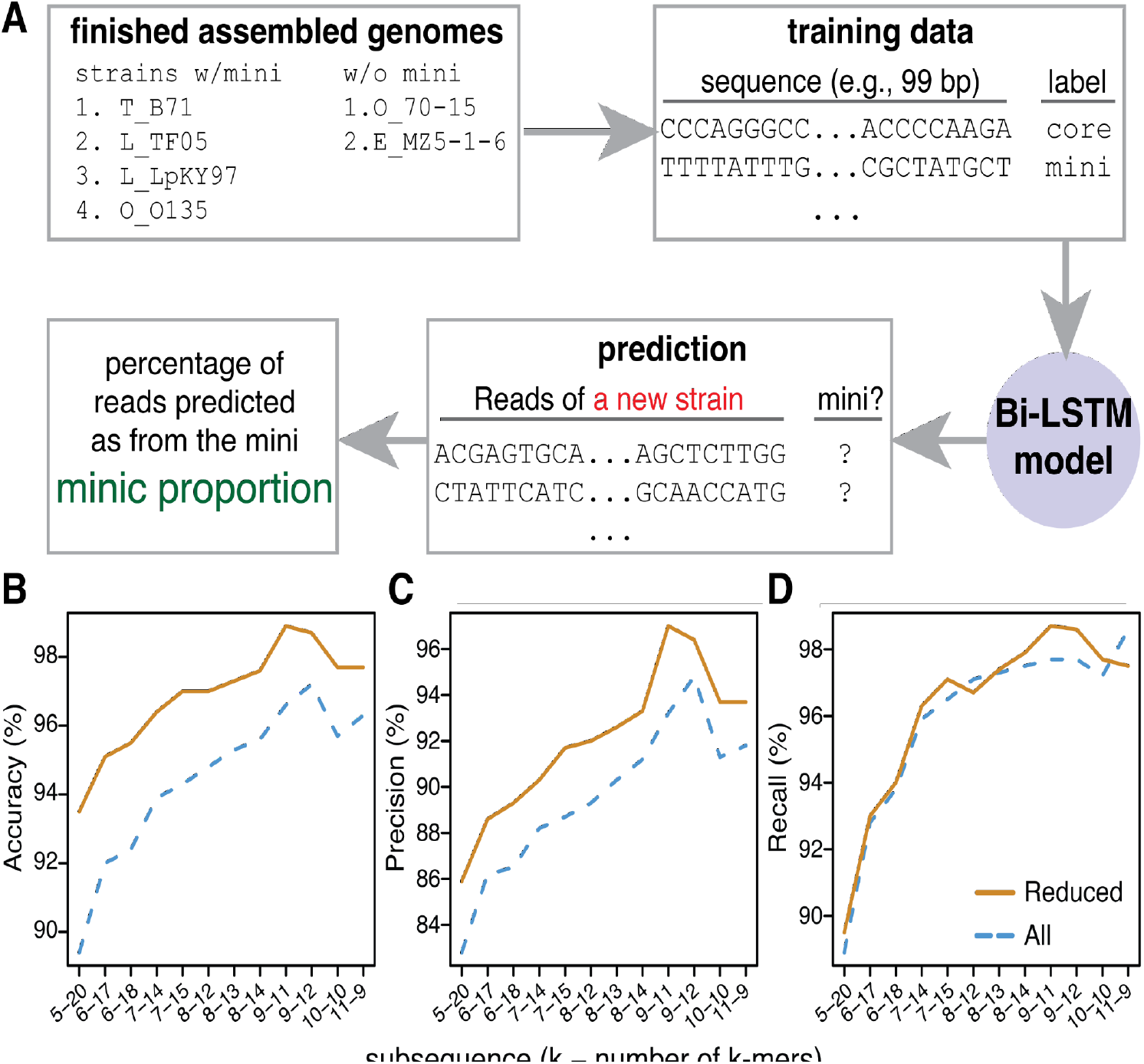
Overview of predicting sequences from mini-chromosomes. (**A**) Finished assembled genomes of strains with or without the mini-chromosome were used to generate training data. The training data include DNA fragments with the length around 100 bp and labeled with the origins from either core-or mini-chromosomes. The deep learning model was trained and the optimal model was selected for predicting the origin of each sequencing read from a new strain. The miniC proportion, which is the percentage of reads predicted to originate from mini-chromosomes, is the value for inference of the presence of the mini-chromosome in the strain. (**B**-**D**) Average performance metrics for models trained using different subsequences, each of which consists of multiple k-mers. Each x-axis label specifies the size of k and the number of k-mers (e.g., 5-20 stands for a 100 bp subsequence with 20 5-mers). Performance was evaluated for all genomic data (All, blue lines) or after removing common sequences shared between core- and mini-chromosomes (Reduced, orange dashed lines).

### Training of Bi-LSTM models

Short sequences (e.g., 99 bp) extracted from the six mini-chromosomes and 42 core-chromosomes were termed subsequences (**Figure 1A**). Each subsequence was then tokenized into non-overlapping k-mers (e.g., 9-mer). Afterwards, the tokenized data were split into train, validation, and test sets with a 80/10/10 split. Models were trained on the train set and evaluated for training performance and hyperparameters selection using the validation set. The selected models were finally evaluated using the test set. When constructing the training dataset, we encountered imbalanced training sequence data from core- and mini-chromosomes in which the total length of core-chromosomes was markedly larger than the total length of mini-chromosomes (**Table 1**). To create balanced training data from core- and mini-chromosomes, we extracted subsequences with the step size of 1 bp from mini-chromosomes and the step size of 27 bp from core-chromosomes.

Models were trained with DNA sequence data of different k-mer sizes, ranging from 5 to 11 mers, and subsequence lengths. Lengths of subsequences were limited to around 100 bp because lengths of whole genome sequencing (WGS) data, or reads, of most *M. oryzae* strains to be used for the prediction are around 100-150 bp. Overall, the evaluation on the validation data showed that models trained with the 9-mer attained the highest scores in both accuracy and precision, and the recall score was close to the highest score achieved by using 11-mer (**Figure 1B-D, Table S1**). The assessment with the models on the test data set showed the consistent evaluation result (**Table S2**). We previously showed common sequences, particularly transposable elements, occurred in core- and mini-chromosomes (Peng *et al*., 2019), which created ambiguous sequence examples that did not have a clear class distinction. To examine if the occurrence of these common sequences impacted the model, we re-trained models using genomic data where common sequences were identified per strain and removed from the training data. Model performance on the validation data was improved when removing these sequences (**Figure 1B-D**), and the model trained using 9-mers became the best for accuracy, precision, and recall (**Table S3**). Within the 9-mer model, the two subsequences lengths of 99 bp and 108 bp did vary for model performance, and the model trained using the 99 bp subsequences (nine 9-mers) attained better scores: 98.9% accuracy, 97.0% precision, and 98.7% recall on both the validation and test datasets (**Figure S1, Table S3, S4**). This model was used for subsequent analysis.

### Survey of presences of mini-chromosomes in cereal blast strains

The optimized Bi-LSTM model was used to examine the presence of a mini-chromosome in *M. oryzae* isolates whose WGS data were available. The probability of the mini-chromosome origin of each WGS read was estimated, and reads with the prediction probability larger than 0.99 were classified as mini-chromosome reads (**Figure S2**). The proportion of mini-chromosome reads among all examined reads, referred to the miniC proportion, was determined for each *M. oryzae* isolate. In total, WGS data of 252 *M. oryzae* isolates from multiple pathotypes were analyzed, resulting in miniC proportions ranging from 0.7% to 9.3% (**Figure 2A, Table S5**). Three isolates, B71 (MoT), P3 (MoT), and LpKY97 (MoL), carrying at least one mini-chromosome had miniC proportions of 3.5%, 5.8%, and 5.6%, respectively. Note that the P3 and LpKY97 genome contained two mini-chromosomes based on the previous reports (Peng *et al*., 2019; Rahnama *et al*., 2021b). In contrast, the miniC proportions of four isolates with no mini-chromosomes, 70-15,Guy11 (MoO), MZ5-1-6 (MoE), T25 (MoT), were 0.8%, 0.9%, 1.1%, and 0.9%, respectively. Based on miniC proportions of these isolates, we used 1.5% as the miniC proportion threshold to classify isolates with or without the mini-chromosome.

**Figure 2.**
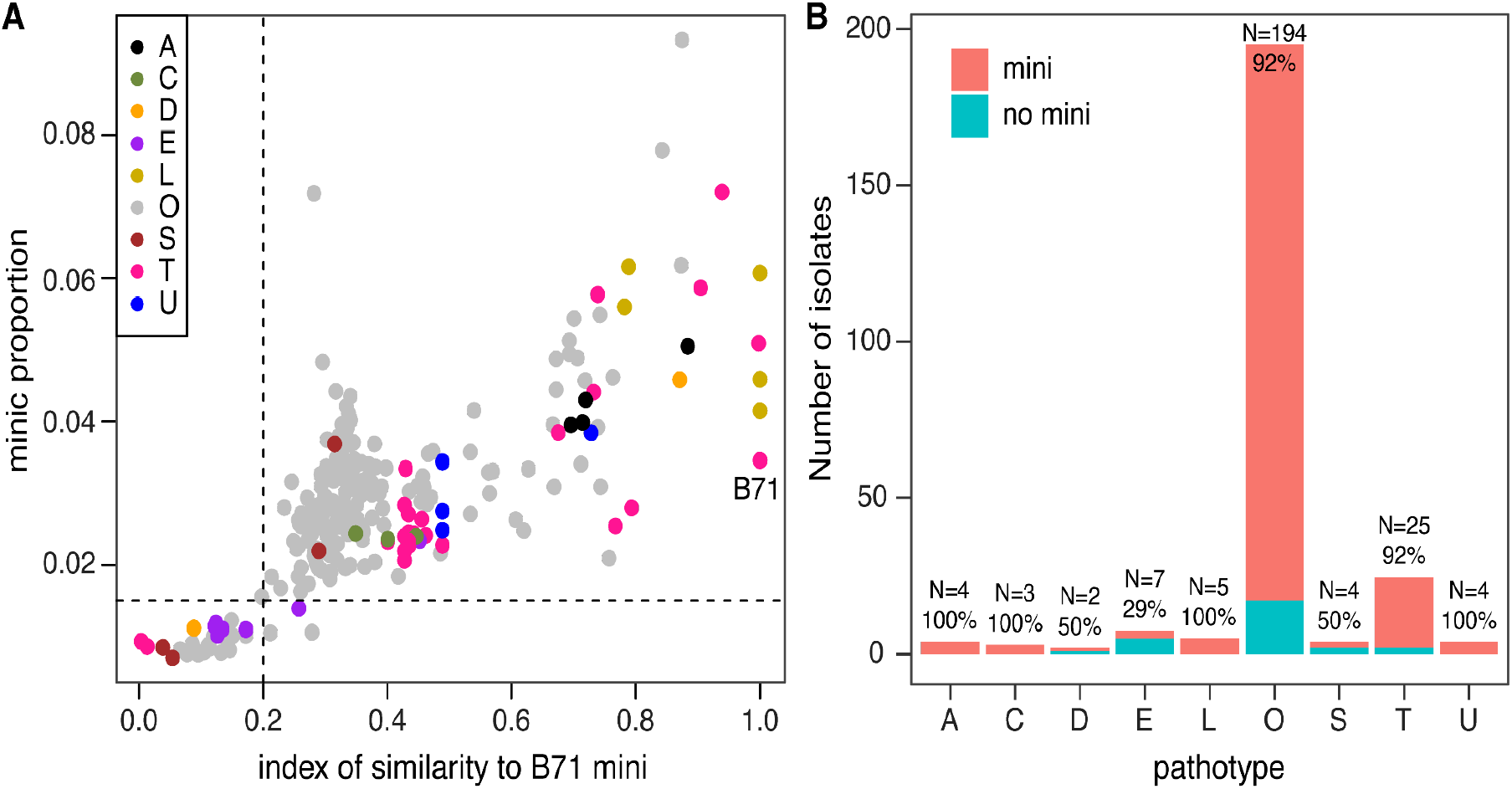
The presence of the mini-chromosome in isolates of diverse pathotypes. (**A**) The miniC proportion of each isolate was estimated using whole genome sequencing reads. The index of similarity to the B71 mini-chromosome represents the proportion of the B71 mini-chromosome that was not detected as deletion genomic regions using the CGRD pipeline. Dash lines signify the miniC proportion threshold of 1.5% and the similarity index threshold of 0.2 used to determine if an isolate carried the mini-chromosome. Letters stand for host species on which the strains were isolated in the field (e.g., A=*Avena*; C=*Cenchrus*; D=*Digitaria*; E=*Eleusine*; L=*Lolium*; O=*Oryza*; S=*Setaria*; T=*Triticum* and U=*Urochloa*). (**B**). Distribution of the number of isolates with and without the mini-chromosome in each pathotype. Total numbers of isolates and percentages of isolates with the mini-chromosome are labeled on top of bars.

The Comparative Genomics Read Depth (CGRD) pipeline was employed to identify the genomic regions of the B71 mini-chromosome that were absent in each isolate (Peng *et al*., 2019; Lin *et al*., 2021). The proportion of the B71 mini-chromosome that was not detected as absence regions represents the similarity of the potential mini-chromosome of an isolate to the B71 mini-chromosome, which was referred to as the index of similarity to the B71 mini-chromosome. Index values of 252 strains ranged from 0.03 to 1. A higher index of similarity indicates a higher possibility that an isolate carries the mini-chromosome. Based on the index values of isolates known to carry at least a mini-chromosome or none, the index threshold of 0.2 was used to classify isolates with or without the mini-chromosome.

Comparison between the prediction result from the Bi-LSTM model with the result using the CGRD approach showed that the two methods were highly similar. Specifically, the prediction of the mini-chromosome presence in 98.4% (248/252) isolates were the same. In total, 223 were predicted to contain the mini-chromosome using both approaches, indicative of a substantial presence of mini-chromosomes across *M. oryzae* strains. The results also indicated that different pathotypes had varying levels in mini-chromosome prevalence (**Figure 2B**). More than 90% of both 195 MoO and 25 MoT isolates were predicted to carry mini-chromosomes. All isolates collected from *Avena* spp., *Cenchrus* spp., *Lolium* spp., and *Urochloa* species, and half of isolates from *Digitaria* spp., and *Setaria* spp., were predicted to contain mini-chromosomes. The mini-chromosome was the least prevalent in pathotype MoE, of which 29% (2/7) isolates were predicted to contain mini-chromosomes. Note that the number of isolates of each of the pathotypes other than MoO and MoT is relatively small, ranging from 2 to 7. Four MoO isolates, namely IR0088, IR0095, JP0091, and YN8773-19, proved difficult to predict and produced different predictions from the two prediction approaches. The miniC proportions of the four isolates were from 1.3% to 2.2%, while the indexes of similarity to the B71 mini-chromosome were from 0.15 to 0.28. Both predictions of all the four strains were close to the respective thresholds.

### Prediction using subsets of reads

To determine the minimal amount of sequencing reads required for reliable prediction, four isolates with known numbers of mini-chromosomes were selected for a simulation. These four isolates included P3 with two mini-chromosome, B71 with one mini-chromosome, and two mini-chromosome-free isolates: T25 and Guy11 (Peng *et al*., 2019; Rowe *et al*., 2023). Random reads, from 1,000 to 300,000, were subsampled from the forward read sets of the original paired-end WGS reads, with subsampling repeated five times per isolate. As expected, the variation of miniC proportions was higher when a low amount of reads were used for the prediction (**Figure 3**). The simulation from all the four isolates consistently showed that the prediction of the miniC proportion was not very reliable when the number of reads was less than 20,000. However, even when using a low number of reads, the predicted proportion value of mini-chromosomes did not deviate dramatically from the prediction value obtained using the original full read set. When 50,000 or more reads were used, the predicted miniC proportion was very close to that using the original read set, which included millions of reads. Based on the simulation result, 100,000 and more reads are conservatively recommended for an accurate prediction of miniC proportions using our Bi-LSTM model.

**Figure 3.**
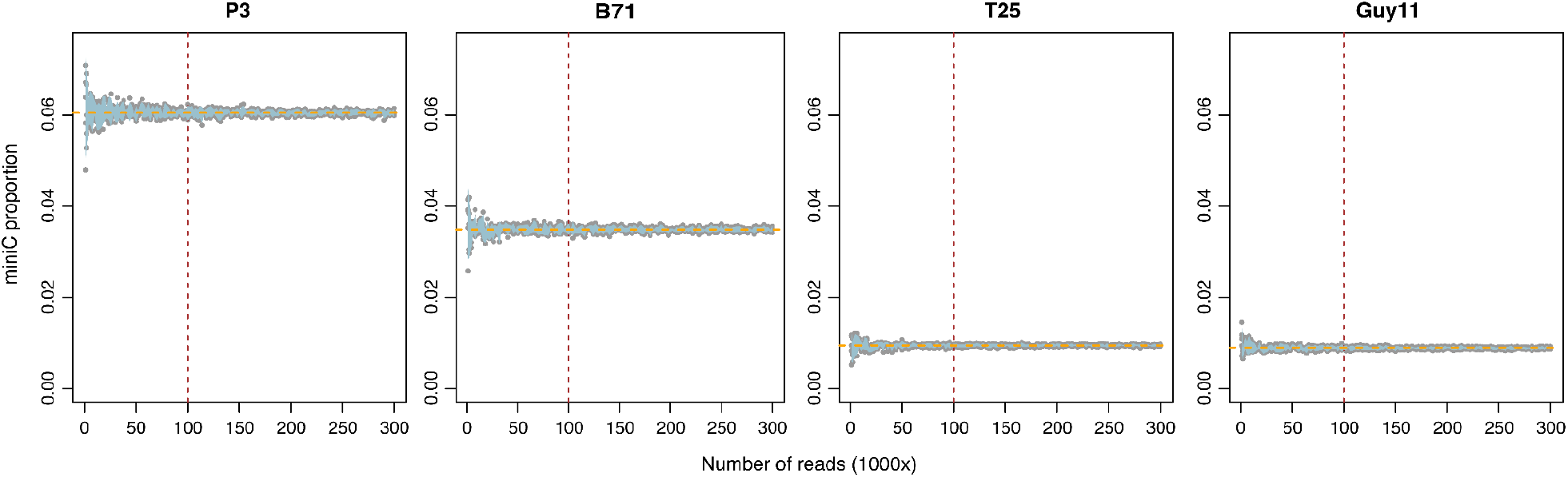
Prediction of miniC proportions using subsampled reads. MiniC proportions (Y-axis) from five times of simulations for each of four isolates (P3, B71, T25, and Guy11) were plotted versus numbers of reads used (X-axis). Each gray dot represents a predicted miniC proportion using a certain number of reads randomly extracted from an original read set of the corresponding isolate. The light blue shades were plotted using 95% confidence intervals from five simulations. Orange horizontal dash lines indicate the predicted miniC proportion values using the original full reads. Brown vertical dash lines point the recommended minimum read number for prediction.

### Applications to identify mini-chromosome-associated sequences

In addition to predicting if a strain contains a mini-chromosomes, we applied the Bi-LSTM model to predict if a DNA sequence from an assembly (termed contig hereafter) represented a mini-chromosome. We split each contig into continuous 99 subsequences and classified each to either core-or mini-chromosome based on the model prediction. The proportion of mini-chromosome subsequence of a contig, referred to the miniC proportion of a contig, indicates the extent to which the contig shares similarity to the mini-chromosome. To test the prediction strategy, the genome assembly of B71, including seven core-chromosomes and one mini-chromosome, was subjected to the analysis. As a result, the miniC proportion of the mini-chromosome was 54.8%, which was markedly higher than miniC proportions of core-chromosomes ranging from 0.1% to 1.7%, (**Figure 4A, Table S6**). A previous study produced draft genome assemblies for MoO FR13 **(Figure 4B)**, MoS US71 **(Figure 4C)**, MoE CD156 **(Figure 4D)**, and demonstrated that all three contained mini-chromosomes by CHEF analysis (Langner *et al*., 2021). MiniC proportions of contigs from each drafted assembly were determined and used to infer if a contig was derived from the mini-chromosome. Eight contigs larger than 100 kb previously found to be mini-chromosome derived were supported by the miniC proportion data, which identified additional five contigs (**Table S6**). In the same study, MoT BR32 was found to contain no mini-chromosomes. Consistently, the miniC proportions of all contigs are small, ranging from 0.5% to 3.1% **(Figure 4E, Table S6)**. Furthermore, we assembled a new MoT genome T3 into seven chromosomes, indicative of no mini-chromosomes. All these seven chromosomes had small miniC proportions (0.5-1.4%) and can be assigned to core-chromosomes 1 to 7 **(Figure 4F, Table S6)**. Collectively, the Bi-LSTM model we constructed can be used to differentiate contigs belonging to core-or mini-chromosomes.

**Figure 4.**
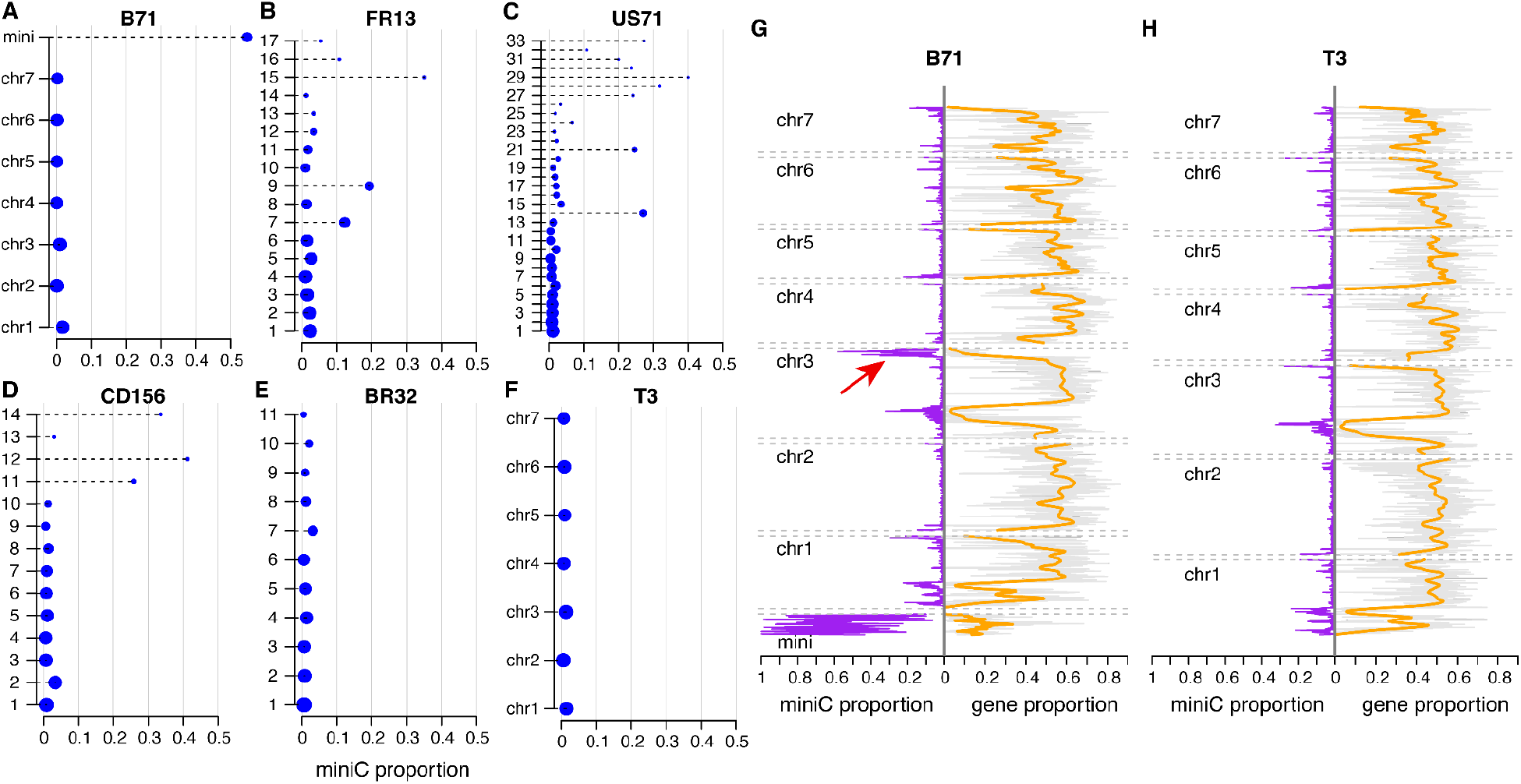
Prediction of miniC proportions in assembled contigs or chromosomes. (**A**-**F**) Assembled contigs or chromosomes of six strains, including five strains that are not included in the training data, were subjected to the prediction. The miniC proportion represents the proportion of sequences in each contig/chromosome predicted as miniC sequences. Y-axes signify names of contigs/chromosomes, which are listed on the column of “contig_id” in **Table S6**. The same rule is applicable for other contig names. Sizes of blue dots indicate the bp length of contigs/chromosomes. (**G, H**) miniC proportions per 30-kb interval (purple) and proportions of genic sequences per interval along each chromosome (gray). Orange curves represent LOWESS estimates of genic proportions. The red arrow indicates a core-chromosome region with a high miniC proportion value. B71 genome version: B71Ref2; T3 genome version: T3v1.

To scan along individual chromosomes, the calculated miniC proportions were determined for 30 kb intervals of each chromosome of B71 and T3, which carried one and zero mini-chromosomes, respectively. Almost all intervals of the B71 mini-chromosome had a miniC proportion larger than 10%. Many intervals on the ends of core-chromosomes showed a relatively high miniC proportion, indicating that they possess sequence features associated with the mini-chromosome. Notably, a region at the end of B71 chromosome 3 contained sequences with a miniC proportion level similar to the mini-chromosome. This region is absent in the genome of T3, which did not contain the mini-chromosome (**Figure 4G and 4H**). The region represents a potential translocation event from the mini-chromosome to the core-chromosome.

## DISCUSSION

In this study, we employed a Recurrent Neural Network (RNN) deep learning technique, specifically a Bi-directional Long Short-Term Memory (Bi-LSTM) network, to model the origin of DNA sequences as belonging to core-or mini-chromosomes. The optimized Bi-LSTM model enables examination of the core-or mini-chromosome origin using the input data of WGS reads, assembled contigs or chromosomes, and DNA sequence fragments. The model was trained using multiple genomes with or without the mini-chromosome, learning genomic features from divergent core- and mini-chromosomes. As compared to alignment-based approaches such a CGRD, the Bi-LSTM model has reduced reference genome bias because it was trained with multiple divergent genomes from different host-adapted pathotypes (MoO, MoT, MoL and MoE) and with differing mini-chromosome composition. Also, the Bi-LSTM model is able to analyze both non-repetitive and repetitive sequences, overcoming a common problem of repetitive sequences limiting whole genome analysis. We show that Bi-LSTM prediction is accurate and reliable even using a small amount of WGS read data. Crosstalk between core-and mini-chromosomes in *M. oryzae* was previously hypothesized (Peng *et al*., 2019). Our efforts to scan assembled chromosomes identified the end of chromosome 3 in the B71 as being highly similar to the mini-chromosome. Given that this region is absent in the B71 related strain, T3, this region may represent a genome structural variation arising from a translocation event from the mini-chromosome. Future analysis of more high-quality reference level assemblies of more diverse *M. oryzae* strains will further illuminate potential core genome variation influenced by the mini-chromosome.

Analysis of 252 *M. oryzae* isolates reveals the prevalence of mini-chromosomes in at least some field isolates of all *M. oryzae* host-adapted pathotypes that we investigated. Specifically, 92% of 196 rice isolates were predicted to carry the mini-chromosome. The result is consistent with a previous examination of mini-chromosomes conducted using electrophoretic karyotyping, which found 93% of 14 rice isolates harbored the mini-chromosome (Orbach *et al*., 1996). In the same study, none of seven wheat isolates carried the mini-chromosome. However, our analysis showed that 92% of wheat strains carried the mini-chromosome. The discrepancy appears to be related to the isolation period for these wheat isolates relative to the first report of wheat blast disease in 1985 in Brazil (Igarashi, 1986). All isolates (T1 to T7) examined in Orbach et al. (1996) were early wheat strains collected in 1988 or earlier. From our analyses, none of the three early wheat strains (T3, T25, and BR32 collected in 1986, 1988, and 1991, respectively) carried mini-chromosomes. In contrast, all wheat isolates collected after 2005 carried the mini-chromosome. The result indicated that wheat strains with the mini-chromosome were preferentially selected in the field over the period from 1990s. All the Lolium isolates examined, which are closely related to wheat isolates, carried the mini-chromosome. Among all pathotypes analyzed, strains of *Eleusine* (MoE) pathotype generally lacked mini-chromosomes. Combined with two different MoE strains analyzed in the Orbach *et al*. study, 78% of MoE (7/9) isolates contained no mini-chromosomes. Orbach *et al*. (1996) reported that all 18 fully fertile field isolates and derived fertile laboratory strains, which are able to serve as a female and produce perithecia in sexual crosses with strains of opposite mating type, lacked mini-chromosomes. They also showed that mini-chromosomes fail to segregate normally in sexual crosses. In contrast, mini-chromosomes occur frequently in strains that lack sexual fertility. Our results support the correlation between lack of mini-chromosomes and female fertility in sexual crosses since MoE strains, in general, are known to possess high levels of sexual fertility (Valent *et al*., 2020) and they generally lack mini-chromosomes. Also in support of this correlation, recent data shows that both the *Triticum* and *Lolium* pathotypes co-evolved in a recent multi-hybrid sexual swarm that resulted in the re-partitioning of standing variation present in at least eight individuals from 5 different host-adapted pathotypes, including the *Eleusine* pathotype (Inoue *et al*., 2017; Rahnama *et al*., 2021a). After emergence of populations adapted to wheat and to *Lolium* spp., asexual reproduction systems seem to currently predominate during infections in the field, perhaps allowing mini-chromosome accumulation (Maciel *et al*., 2014). Although current data suggests a correlation, further studies are needed to determine the precise role of the mini-chromosome in sexual fertility. Our mini-chromosome prediction model provides a new tool for addressing the question and tracking mini-chromosome presence in evolving populations of the blast fungus.

Our prediction model can be further improved by training using additional core- and mini-chromosome sequencing data. In particular, more mini-chromosome sequencing data will allow capturing the high-level diversity among mini-chromosomes. The sampling of *M. oryzae* isolates is heavily skewed to rice and wheat infecting lines, and greater sampling of more diverse pathotypes will also improve this approach. In addition to the prediction of the presence of the mini-chromosome, the model may predict the number of mini-chromosomes in each isolate. Nevertheless, this study demonstrates the potential of deep learning techniques in genomics for predicting the presence of specific genomic elements. We anticipate that in the near future, using improved explainable deep learning techniques, the critical sequence components of mini-chromosome DNAs may be identified by learning from massive genomic data to further understand the origin and evolution of mini-chromosomes.

## METHODS

### Near-finished genome assemblies for model training

Near-finished genome assemblies of isolates that are known to carry or not to carry the mini-chromosome were collected for training Bi-LSTM models, which include the assemblies from mini-chromosome-bearing isolates B71 (MoT, Genbank accession: GCA_004785725.2) (Peng *et al*., 2019; Liu *et al*., 2022), LpKY97 (MoL, Genbank accession: GCA_012272995.1) (Rahnama *et al*., 2021b), TF05-1 (MoL, Genbank accession: GCA_002924755.1), and O135 (MoO, Bioproject accession: PRJNA1006359) (Rowe *et al*., 2023), as well as assemblies from mini-chromosome-free isolates 70-15 (MoO, Genbank accession: GCA_000002495.2) (Dean *et al*., 2005) and MZ5-1-6 (MoE, Genbank accession: GCA_004346965.1) (Gómez Luciano *et al*., 2019).

### Identification common sequences between core- and mini-chromosomes

Alignment was performed between core- and mini-chromosomes between each of B71, O135, LpKY97, and TF05-1 with NUCmer (Marçais *et al*., 2018). Alignments with at least 105 bp matches and 95% identity were retained. Alignment regions were merged if neighboring alignments were within a 100 bp distance and sequences were extracted from both core- and mini-chromosomes. All common sequences identified from these four genomes and the mitochondrial sequence of B71 were combined to form a database of sequences excluded from the training.

### Bi-LSTM models

The Bi-LSTM model was implemented using Python with TensorFlow and Keras libraries. The architecture consisted of an input layer, hidden LSTM cells layers, and an output layer. The input layer consists of an embedding layer that encodes the input tokens (such as the 11 9-mers tokens of the 99 bp sequence) into vectors of size 128. The hidden layers consisted of two Bidirectional LSTM layers, each with 256 hidden units, stacked on top of each other with the hyperbolic tangent (tanh) activation function. The output of the last Bi-LSTM hidden layer was connected to a dense output layer with sigmoid activation function. A model for 99b bp sequence with 11 9-mer tokens contained a total of 34,212,225 trainable parameters. The selection of the model architecture and hyperparameters was informed by experimenting with a range of values and selecting those that resulted in the best performance on the validation set.

For training the Bi-LSTM model, backpropagation through time (BPTT) was employed, using binary cross entropy loss as the loss function. The optimization was performed using Adam optimizer with learning rate of 0.001. The training and validation data sets, such as sequences with 99 bp with 11 9-mers and labeled with either “core” or “mini”, were encoded using one-hot encoding and used to train and evaluate the model. A mini-batch size of 2048 was used for the training and evaluation at each epoch. To optimize the training and prevent overfitting, an early stopping criteria based on validation loss was implemented. The model was trained for a maximum of 150 epochs with patience of 15 epochs. If the validation loss did not improve over the subsequent 15 consecutive epochs, the training was stopped. The model weights corresponding to the lowest validation loss were restored, representing the best-performing model. Subsequently, the final trained model was then tested on an independent test dataset to evaluate its overall performance.

### Draft genome assemblies of isolates with and without mini-chromosomes

Genome assembly drafts were downloaded from Genbank: GCA_900474655.3 of FR13 (MoO from rice), GCA_900474175.3 of US71 (MoS from foxtail millet), GCA_900474475.3 of CD156 (MoE from goosegrass), and GCA_900474545.3 of BR32 (MoT from wheat). Field isolates FR13, US71, and CD156 all carried the mini-chromosome and BR32 did not contain a mini-chromosome (Langner *et al*., 2021).

### Illumina WGS short-reads of 252 *M. oryzae* isolates

WGS reads were downloaded from Sequence Read Archive (SRA). Data of 252 accessions were collected (**Table S7**). Reads were trimmed with software Trimmomatic (Bolger *et al*., 2014) prior to further analyses.

### Subsampled reads for determining miniC proportions

Random seeds were set for sampling reads from the forward reads of the original paired-end Illumina reads of the isolates of P3, B71, T25, and Guy11. Subsampling was implemented using seqtk (version 1.2). Subsampled reads were then used for the prediction with the optimized Bi-LSTM model.

### Indexes of similarity to the B71 mini-chromosome

WGS reads of each strain were used to compare with WGS reads of B71 to infer the genomic regions of the B71 mini-chromosome that were absent in the isolate through Comparative Genomics Read Depth (CGRD) (Peng *et al*., 2019; Lin *et al*., 2021). The proportion of the B71 mini-chromosome that were not detected as absence regions represents the portion of sequences in the isolate similar to the B71 mini-chromosome, referred to as the index of similarity to the B71 mini-chromosome. A low value of the index indicates the absence of the mini-chromosome.

### Genome sequencing and assembly of early MoT strain T3

The MoT strain T3 was cultured on oatmeal agar (OMA) plates followed by liquid culture under Biosafety Level 3 (BSL3) laboratory in the Biosecurity Research Institute (BRI) at Kansas State University in Manhattan, KS (Valent *et al*., 1986; Liu *et al*., 2022). The detailed procedure for genomic DNA extraction was previously described (Liu *et al*., 2022). Briefly, mycelial mats were collected, lyophilized, and ground for DNA extraction with a CTAB approach. DNA was stored in the TE buffer containing 1 mg/ml RNase. Approximately 50x 2×150 bp Illumina data were produced at Novogene USA. Nanopore long reads were generated using the same genomic DNAs per the procedure described previously (Liu *et al*., 2022). The genomic DNA was subjected to a size selection (>20 kb) using BluePippin Gel Cassette (Sage Science, USA, Cat.# BLF7510), followed by a library construction using the SQK-LSK110 kit and sequencing using a R9.4.1 flow cell on a MinION Mk1B device (Oxford Nanopore, UK). Nanopore raw FAST5 data were converted to FASTQ reads using the Guppy Basecaller (version 6.3.2). Reads were assembled with Canu (version 2.2) with the parameters of “genomeSize=45m minReadLength=10000 minOverlapLength=1000 correctedErrorRate=0.08 rawErrorRate=0.3 corOutCoverage=60” (Koren *et al*., 2017). The contigs in Canu assemblies were aligned to B71Ref2 to determine the chromosome number and the orientation using NUCmer with the parameters of “-L10000 -I 90”(Marçais *et al*., 2018).

The resulting assembly was polished using Nanopolish (version 0.14.0) and then using Pilon (version 1.24) with Illumina reads (Walker *et al*., 2014; Loman *et al*., 2015).

## Acknowledgements

We thank funding provided by the USDA NIFA award (2018-67013-28511) to S. Liu; USDA NIFA award (2021-68013-33719) to B. Valent; and USDA NIFA award (2021-67013-35724) to S. Liu, B. Valent, D. Cook, and D. Koo; the NSF awards (1741090 and 2311738) to S. Liu; and NSF award (2011500) to S. Liu, B. Valent, D. Cook, and D. Koo. This is contribution no. 24-030-J from the Kansas Agricultural Experiment Station, Manhattan, Kansas.

## Data availability

Nanopore genomic sequencing data of T3 have been deposited in the Sequence Read Archive (SRA) database under accessions PRJNA1002604, and Illumina sequencing data in PRJNA1002398. Related scripts are available at GitHub (https://github.com/PlantG3/miniC).

## Author contribution

NG and SL conceptualized experiments; GL, JH, RB, LCD, HZ, GC conducted experiments; NG, YH, and SL analyzed data; NG, D Caragea, D Cook, BV, and SL wrote the manuscript; all authors reviewed and revised the manuscript.

