## Supplementary Figures for "Using recurrent neural networks to detect supernumerary chromosomes in fungal strains causing blast diseases"

Gyawali et al.

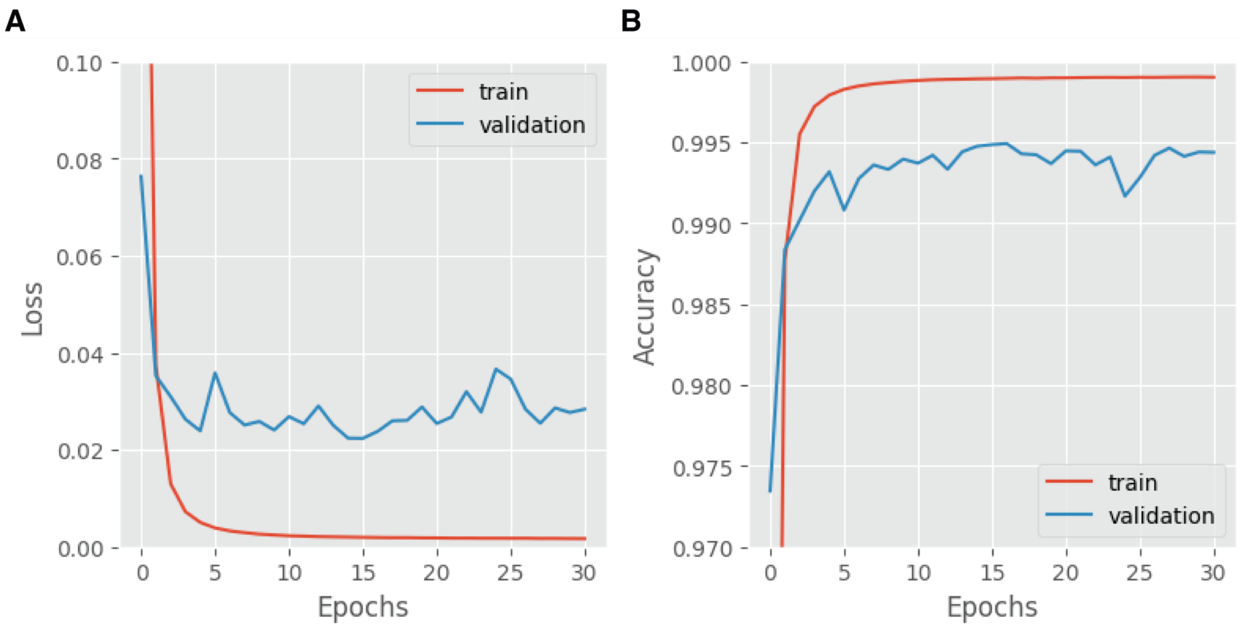


**Figure S1. Prediction loss and accuracy of models during training**
Plot of (**A**) Loss and (**B**) Accuracy at the end of each epoch during the training of the model with 99 bp sequences each of which consists of eleven 9-mer tokens. As shown in (A), the model's performance did not improve after epoch 15 until epoch 30 and the model training stops after 30 epochs. The checkpoint at epoch 15 is restored as the best model.

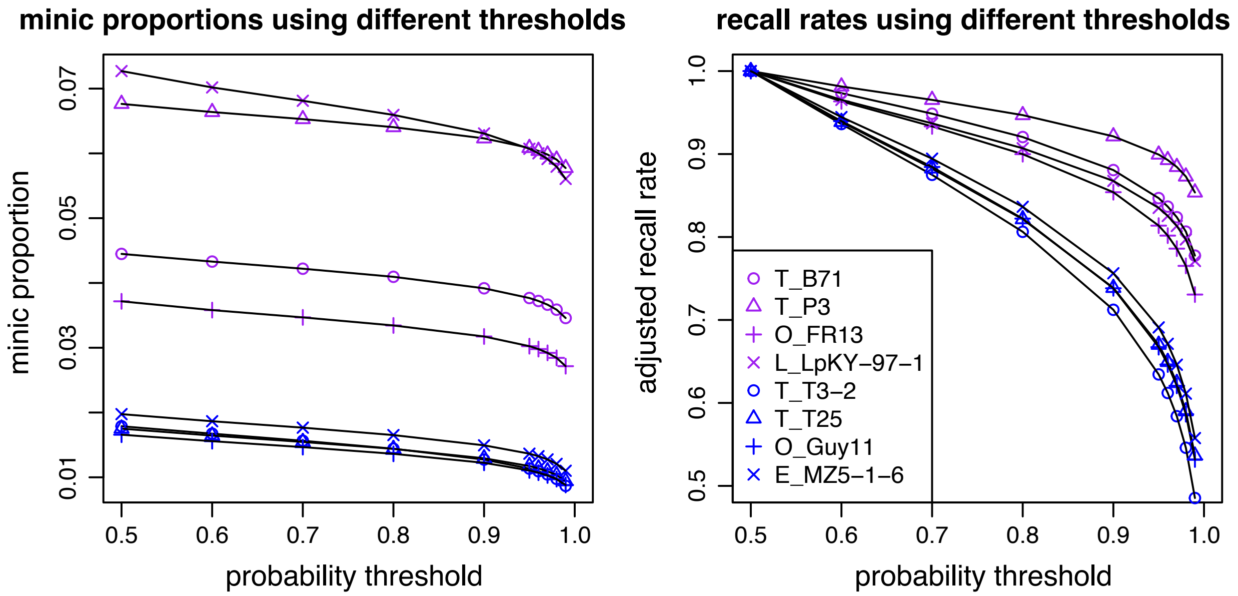


**Figure S2. MiniC proportions and recall rates with different probability thresholds**
By default, the probability of 0.5 was used for the classification. The probability to classify a sequence to be from a mini-chromosome was increased. Plot (**A**) shows the changes of miniC proportions of multiple strains over different probability thresholds; plot (**B**) shows the changes of adjusted recall rates, each of which was the ratio of the number of sequences classified as mini-chromosome sequences using a probability threshold indicated in X-axis to that using the probability threshold of 0.5. Strains indicated in blue had no mini-chromosomes, while strains indicated in purple had at least one mini-chromosome. The recall rates drop faster when the probability threshold increase. Therefore, the probability of 0.99 was finally selected as the threshold for the classification.
